# Modeling statin-induced myopathy with human iPSCs reveals that impaired proteostasis underlies the myotoxicity and is targetable for the prevention

**DOI:** 10.1101/2024.08.23.608904

**Authors:** Xiaolin Zhao, Liyang Ni, Miharu Kubo, Mariko Matsuto, Hidetoshi Sakurai, Makoto Shimizu, Yu Takahashi, Ryuichiro Sato, Yoshio Yamauchi

## Abstract

Statins, HMG-CoA reductase inhibitors, have been widely prescribed to lower circulating low-density lipoprotein cholesterol levels and reduce the risk of cardiovascular disease. Although statins are well tolerated, statin-associated muscle symptoms (SAMS) are the major adverse effect and cause statin intolerance. Therefore, understanding the molecular mechanisms of SAMS and identifying effective strategies for its prevention are of significant clinical importance; however, both remain unclear. Here we establish a model of statin-induced myopathy (SIM) with human induced pluripotent stem cell (hiPSC)-derived myocytes (iPSC-MCs) and investigate the effect of statins on protein homeostasis (proteostasis) that affects skeletal muscle wasting and myotoxicity. We show that treating hiPSC-MCs with statins induces atrophic phenotype and myotoxicity, establishing a hiPSC-based SIM model. We then examine whether statins impair the balance between protein synthesis and degradation. The results show that statins not only suppress protein synthesis but also promote protein degradation by upregulating the expression of the muscle-specific E3 ubiquitin ligase Atrogin-1 in a mevalonate pathway-dependent manner. Mechanistically, blocking the mevalonate pathway inactivates the protein kinase Akt, leading to the inhibition of mTORC1 and GSK3β but the activation of FOXO1. These changes explain the statin-induced impairment in proteostasis. Finally, we show that pharmacological blockage of FOXO1 prevents SIM in hiPSC-MCs, implicating FOXO1 as a key mediator of SIM. Taken together, this study suggests that the mevalonate pathway is critical for maintaining skeletal muscle proteostasis and identifies FOXO1 as a potential target for preventing SIM.

## Introduction

Statins are inhibitors of 3-hydroxy-3-methylgrutaryl (HMG)-CoA reductase (HMGCR), an endoplasmic reticulum (ER) membrane-bound enzyme that converts HMG-CoA into mevalonate (1). HMGCR acts as the rate-limiting enzyme of the mevalonate (MVA) pathway and cholesterol biosynthesis, and its activity is therefore tightly regulated at transcriptional and post-translational levels (2, 3). Mevalonate is then converted into the 5-carbon isoprenoid isopentenyl pyrophosphate (IPP), an essential precursor for synthesizing subsequent isoprenoids, including 10-carbon geranyl pyrophosphate (GPP), 15-carbon farnesyl pyrophosphate (FPP), and 20-carbon geranylgeranyl pyrophosphate (GGPP). Both FPP and GGPP are used for protein prenylation that regulates the function of the modified proteins. FPP and GGPP serve as precursors for dolichol, and for ubiquinone and menaquinone-4 (vitamin K2), respectively. The condensation of two molecules of FPP produces 30-carbon squalene which is then converted to lanosterol and subsequently cholesterol. As such, the MVA pathway is an important metabolic pathway that biosynthesizes not only sterols but also non-sterol isoprenoids. Accordingly, the MVA pathway impacts diverse human diseases (2, 4–6).

Circulating low-density lipoprotein cholesterol (LDL-C) levels positively correlate to the risk of atherosclerotic cardiovascular disease (CVD). Controlling LDL-C levels is therefore among the most promising strategies to reduce CVD risk, and statins have widely been prescribed to lower LDL-C (2, 7, 8). Mechanistically, inhibition of HMGCR by statins suppresses cholesterol biosynthesis in the liver, which leads to the activation of sterol regulatory element-binding proteins (SREBPs) (2). SREBP activation leads to the upregulation of LDL receptor expression in the liver, thereby promoting LDL clearance through receptor-mediated endocytosis (2). Although statins are well-tolerated drugs, observational studies reported that up to 30% of statin takers experience statin-associated muscle symptoms (SAMS), the most common adverse effect of statins (4, 9–11). SAMS includes myalgia (muscle pain), myopathy, and rhabdomyolysis, and rhabdomyolysis is the most severe form of statin-induced myopathy (SIM). Due to the higher frequency of rhabdomyolysis, cerivastatin was withdrawn from the market in 2001. SAMS is caused by the myotoxicity of statins and is the major background of statin intolerance, resulting in the cessation of statin therapy. Satin intolerance increases the risk of CVD in patients who discontinue the treatment, and therefore overcoming SAMS is a clinically important challenge.

Statin myotoxicity suggests that the MVA pathway and MVA-derived metabolites play a critical role in skeletal muscle homeostasis. Genetic evidence supports this notion; skeletal muscle-specific deletion of the *Hmgcr* gene leads to severe myopathy in mice (12). Furthermore, recent studies have discovered that the *HMGCR* gene mutations cause limb-girdle muscular disease (LGMD), an autosomal progressive myopathy, which shares similar phenotypes with SIM (13, 14). In both skeletal muscle-specific *Hmgcr* knockout mice and the LGMD patients with *HMGCR* mutations, the administration of mevalonolactone, a lactone form of MVA, at least partly restores the myopathic phenotypes (12, 13). However, how muscular HMGCR inactivation induces myotoxicity remains incompletely understood, particularly in human myocytes due to their limited availability; only a few studies have used primary human myocytes (15–17). Supplementation of the isoprenoid GGPP, not cholesterol protects C2C12 myocytes from statin myotoxicity, suggesting that depletion of GGPP mediates the myotoxicity (18). Multiple mechanisms of SIM have been proposed based on in vitro and in vivo studies using rodent models (4, 19, 20). These mechanisms include an increase in apoptosis, mitochondrial dysfunction, activation of proteasomal protein degradation, and inhibition of protein synthesis. Some of these are explained by the inactivation of the protein kinase Akt, a master regulator of cell survival and metabolism (21). Protein homeostasis (hereafter referred to as proteostasis) plays a key role in the regulation of skeletal muscle mass, and therefore impaired proteostasis underlies multiple myopathies and muscle wasting (22, 23). However, which Akt downstream effector plays a more direct role in developing SIM remains to be determined.

Human induced pluripotent stem cells (hiPSCs) offer excellent cellular models for investigating mechanisms of various human diseases and for developing and evaluating therapeutic strategies (24, 25). In this work, we establish a SIM model with hiPSC-derived myocytes (hiPSC-MCs) and investigate mechanisms of SIM in human myocytes. Our hiPSC-based SIM model reveals that statins impair proteostasis in a manner dependent on the MVA pathway. The impaired proteostasis by statins involves the inactivation of Akt and dysregulation of its downstream effectors, including mTOR complex 1 (mTORC1), GSK3β, and FOXO1. Furthermore, using hiPSC-MCs, we identify FOXO1 as a potential target for the prevention of SIM.

## Results

### Modeling statin-induced myopathy using hiPSC-MCs

Statins inhibit HMGCR and disrupt the mevalonate (MVA) pathway (**Figure 1A**). We first attempted to establish a SIM model using human iPSC-derived myocytes (hiPSC-MCs) and cerivastatin, a statin with strong myotoxic effects. As expected, cerivastatin treatment induced SREBP-2 processing and increased mRNA levels of SREBP-2 target genes, *HMGCS1* and *HMGCR* in hiPSC-MCs (**Figures 1B, C**). We next examined the effect of cerivastatin on cell toxicity and the diameters of myocytes. The results showed that treating hiPSC-MCs with cerivastatin increases cytotoxicity and decreases cell viability in a dose-dependent manner (**Figure 1D**). The myotoxic effects of cerivastatin were observed at as low as 0.3 μM. We next measured the diameters of myosin heavy chain (MHC)-positive myocytes in the presence of cerivastatin and found that cerivastatin treatment dose-dependently decreases the diameters of myocytes by 25–40% (**Figure 1E, F**). Statin myotoxicity was also observed with other statins (**Figure S1**). These results indicate that hiPSC-MCs serve as a useful model to study statin myotoxicity in vitro.

**Figure 1.**
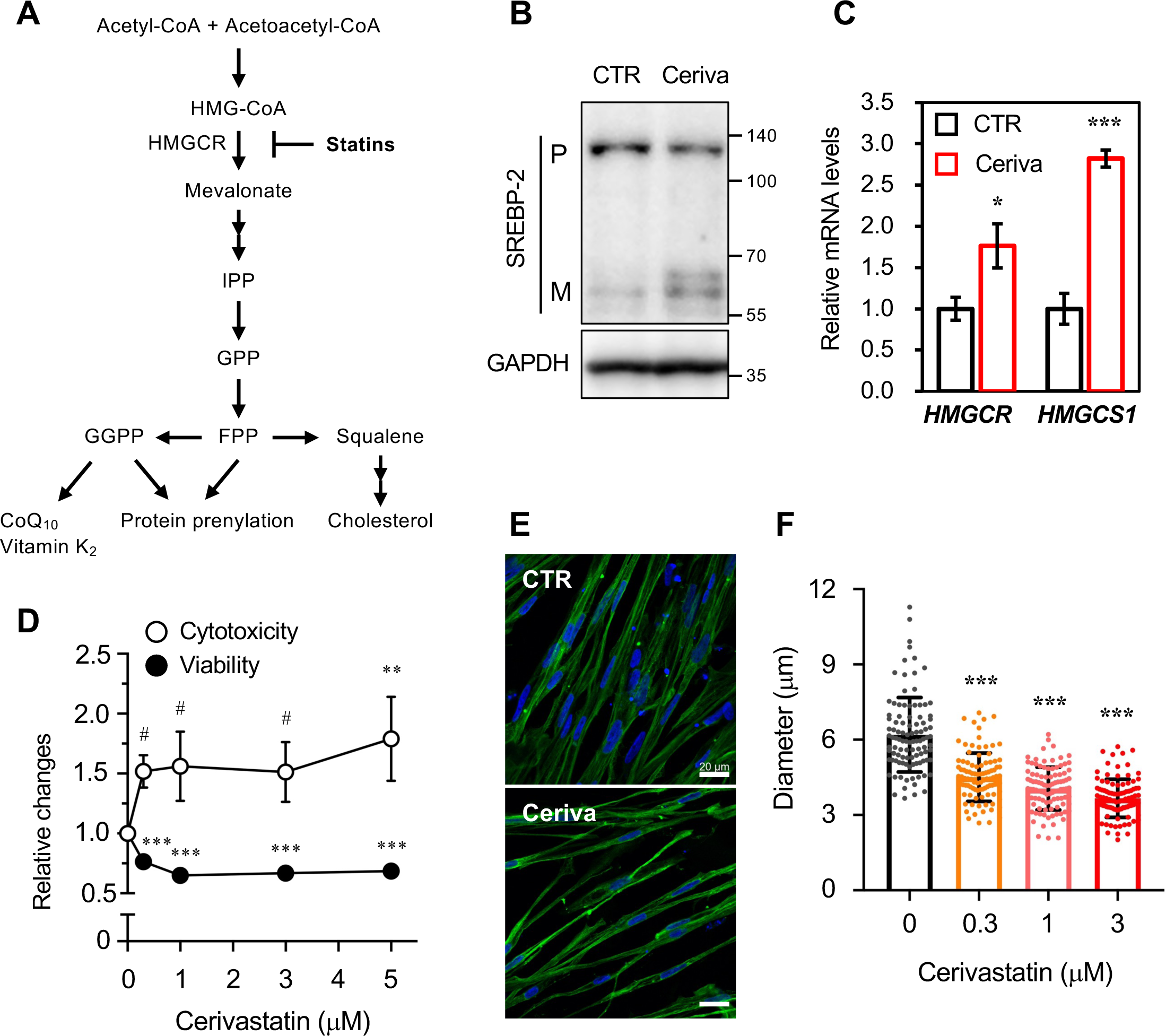
Establishment of statin-induced myopathy with hiPSC-MCs. (A) The MVA pathway. (B) SREBP-2 processing in hiPSC-MCs. hiPSCs (414C2) were differentiated into hiPSC-MCs in 6-well plates. On day 5, hiPSC-MCs were treated with 1 μM cerivastatin for 16 h. Whole cell lysate was subjected to immunoblot analysis, and SREBP2 precursor (P) and mature (M) forms were detected as described in Materials and Methods. GAPDH was detected as an internal control. (C) *HMGCS1* and *HMGCR* mRNA expression in hiPSC-MCs. hiPSC-MCs (414C2) were treated with cerivastatin as above. *HMGCS1* and *HMGCR* mRNA levels were assessed by qRT-PCR. Data represent mean ± SD (n=3). Statistical analysis was performed by student’s *t*-test (* p < 0.05, *** p < 0.001). (D) Effect of cerivastatin on cell viability and cytotoxicity in hiPSC-MCs. hiPSC-MCs (414C2) were treated with cerivastatin for 16 h at the indicated concentrations. Cell viability and cytotoxicity were assayed as described in Materials and Methods. Data represent means ± SD (n=3). Statistical analysis was performed by one-way ANOVA with Dunnett’s post hoc test (# p < 0.08, * p < 0.05, ** p < 0.01, *** p < 0.001). (E, F) Effect of cerivastatin on muscle atrophy. hiPSC-MCs (414C2) incubated with the indicated concentration of cerivastatin for 16 h were fixed and stained for myosin heavy chain (MHC; green) and the nuclei (blue) (E). Scale bar, 20 μm. Diameters of MHC-positive myocytes were plotted in panel F. The results are mean ± SD (n=100). Statistical analysis was performed by one-way ANOVA with Dunnett’s post hoc test (*** p < 0.0001).

The MVA pathway biosynthesizes a variety of biologically important and essential isoprenoids and sterols (**Figure 1A**). We therefore sought to determine a MVA-derived metabolite(s) that is critical for statin myotoxicity. We first performed an add-back experiment to identify a metabolite(s) that prevents myotoxicity. hiPSC-MCs were incubated with cerivastatin in the presence or absence of a series of MVA pathway metabolites, and cell viability was assessed. As expected, the addition of MVA, a product of HMGCR, fully restored statin myotoxicity (**Figure 2A**), indicating that the myotoxic effect depends on HMGCR inhibition. In addition to MVA, geranylgeraniol (GGOH), which is converted to geranylgeranyl pyrophosphate (GGPP) within cells, protected hiPSC-MCs from statin myotoxicity. On the other hand, neither GOH nor FOH canceled myotoxicity. Moreover, squalene and cholesterol, distal products of the MVA pathway, also had no protective effects, suggesting that sterol deficiency is not the cause of statin myotoxicity. To validate the effect of MVA and GGOH, we assessed whether they prevent statin-induced atrophy in hiPSC-MCs. Consistent with **Figure 2A**, both MVA and GGOH restored statin-induced atrophy (**Figure 2B, C**). We next attempted to measure cellular contents of non-sterol isoprenoids, including geranyl pyrophosphate (GPP), farnesyl pyrophosphate (FPP), and GGPP by LC-MS/MS. The results showed that cerivastatin treatment depletes both GPP and FPP, while MVA supplementation increased these contents in the presence of cerivastatin (**Figure 2D**). Cellular GGPP contents were under the detection limit in our analysis. Collectively, these results demonstrate that GGPP is an essential MVA pathway metabolite to protect human myocytes from statin myotoxicity.

**Figure 2.**
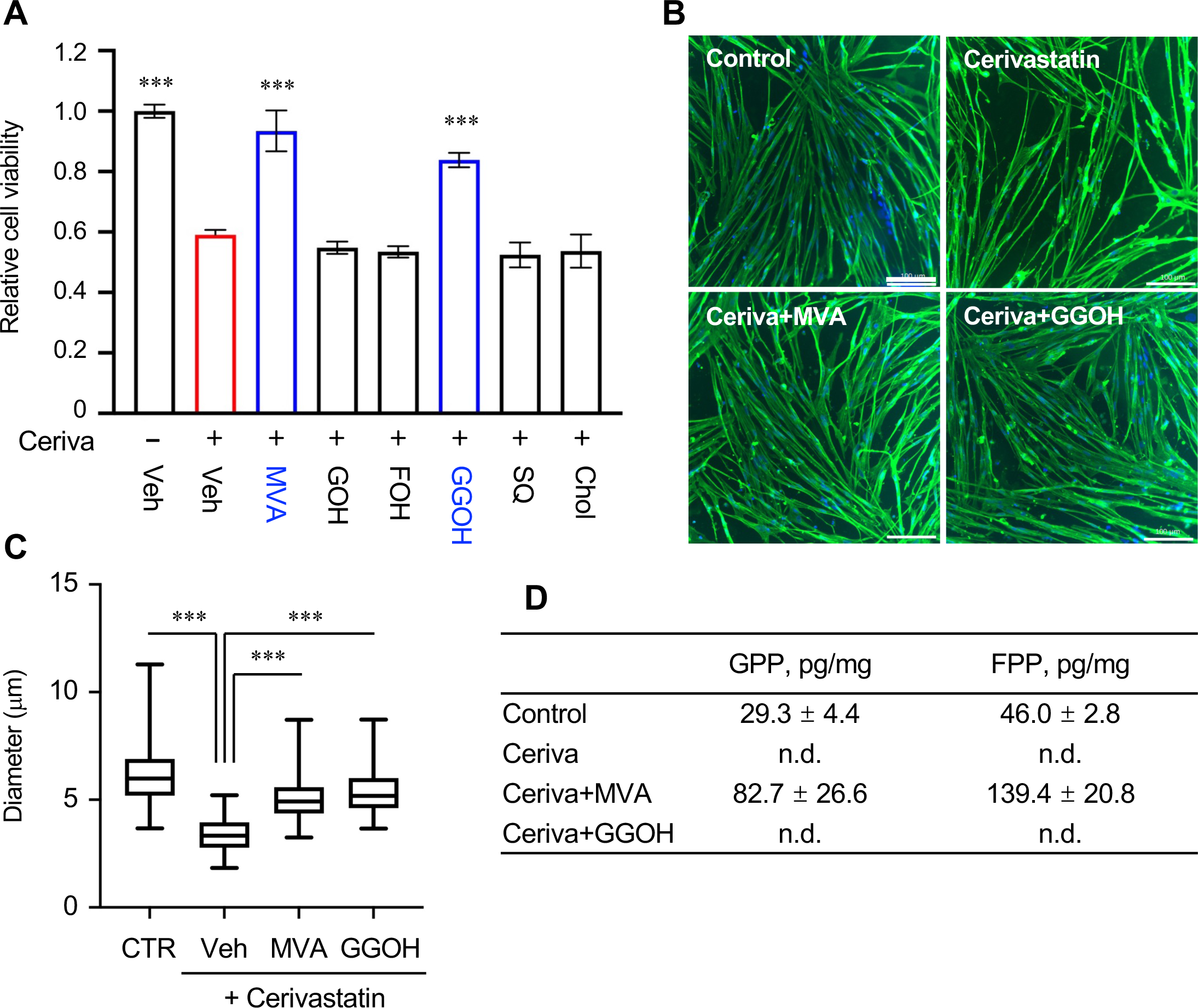
Effects of MVA pathway metabolites on statin myotoxicity in hiPSC-MCs. (A) Effects of MVA pathway metabolites on myocyte viability. hiPSC-MCs (414C2) were incubated with 5 μM cerivastatin in the presence of MVA (200 μM), GOH (100 μM), FOH (100 μM), GGOH (100 μM), squalene (SQ; 20 μM) or cholesterol (Chol; 10 μM) for 16 h. Cell viability was assayed. The results are mean ± SD (n=3). Statistical analysis was performed by one-way ANOVA with Dunnett’s post hoc test by comparing to the cerivastatin-treated group (*** p < 0.0001). (B, C) Effects of MVA and GGOH on muscle atrophy. hiPSC-MCs (414C2) were incubated with 5 μM cerivastatin with or without MVA (200 μM) or GGOH (100 μM) for 16 h. After fixation, hiPSC-MCs were stained for MHC (green) and nuclei (blue) (B). Scale bar, 100 μm. Diameters of MHC-positive myocytes were plotted in panel C. The results are means ± SD (n=100). Statistical analysis was performed by one-way ANOVA with Dunnett’s post hoc test by comparing to the cerivastatin-treated group (*** p < 0.0001). (D) Effect of cerivastatin on cellular isoprenoid contents. hiPSC-MCs (414C2) were incubated with or without 5 μM cerivastatin in the presence or absence of MVA (200 μM) and GGOH (100 μM). Cellular GPP, FPP, and GGPP contents were measured with LC-MS/MS as described in Materials and Methods. Isoprenoid contents were normalized with cellular protein contents. The results are mean ± SD (n=3).

### Statins impair proteostasis in hiPSC-MCs

Proteostasis is crucial for maintaining skeletal muscle mass; muscle atrophy is caused by a reduction in protein synthesis and/or by an increase in protein degradation (22). Therefore, we asked whether statins affect protein synthesis and degradation in hiPSC-MCs. We first examined the rate of de novo protein synthesis using the SUnSET assay which measures the incorporation of puromycin into newly synthesized polypeptides (26). The results showed that cerivastatin reduces de novo protein synthesis in a concentration-dependent manner in both hiPSC414C2- and 409B2-derived myocytes (**Figures 3A, S2A**). Another statin (pitavastatin) also inhibited protein synthesis (**Figure S2B**). We also examined whether the reduction in protein synthesis depends on the MVA pathway. hiPSC-MCs were treated with cerivastatin in the presence or absence of MVA and subjected to the SUnSET assay. The results showed that MVA at least partly restores the statin-dependent reduction of protein synthesis in hiPSC-MCs (**Figure 3B**). These results indicate that the MVA pathway plays an important role in maintaining a capacity for de novo protein synthesis in human myocytes.

**Figure 3.**
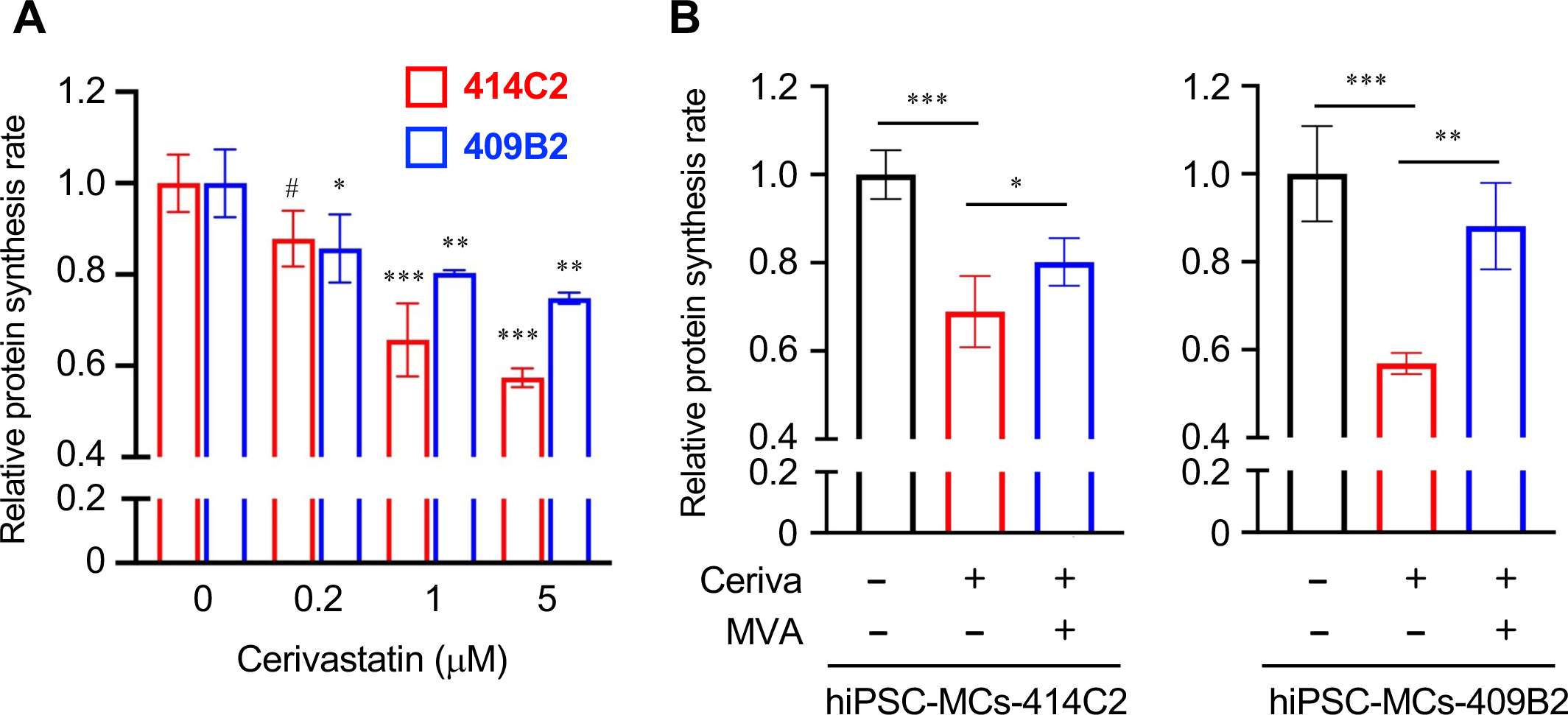
Statins inhibit de novo protein synthesis in hiPSC-MCs. (A) Effects of cerivastatin on protein synthesis. hiPSC-MCs (414C2 and 409B2) were treated with cerivastatin at the indicated concentrations for 16 h. Myocytes were then incubated with puromycin (10 μg/mL) for 1 h. Cell lysate was subjected to immunoblot analysis with anti-puromycin antibody. α-tubulin was used as a loading control. Relative protein synthesis rates are shown (mean ± SD, n=3). The immunoblot images used for the quantification are shown in Figure S2A. Statistical analysis was performed by one-way ANOVA with Dunnett’s post hoc test (# p < 0.1, * p < 0.05, ** p < 0.01, *** p < 0.001). (B) The role of the MVA pathway in protein synthesis. hiPSC-MCs (414C2 and 409B2) were treated with or without 1 μM cerivastatin and/or 200 μM MVA for 16 h, and SUnSET assay was performed. Data represent mean ± SD (n=3). Statistical analysis was performed by one-way ANOVA with Tukey’s post hoc test (* p < 0.05, ** p < 0.01, *** p < 0.001).

Next, we examined whether statins also affect muscle proteolysis in hiPSC-MCs. Atrogin-1 (also known as FBOX32 or MAFbx) and MuRF1 (also known as TRIM63) are two major skeletal muscle-specific E3 ubiquitin ligases that regulate muscle mass (27, 28). We examined the effect of statins on the expression of these two E3 ligases. The results showed that cerivastatin markedly increases Atrogin-1 expression at mRNA and protein levels in hiPSC-MCs (**Figure 4A, B**). In addition to cerivastatin, most statins also upregulated *ATROGIN-1* mRNA levels (**Figure 4A**). The increase in *MuRF1* expression by statins was undetectable or only modest. The upregulation of Atrogin-1 expression by cerivastatin was canceled by the addition of MVA or GGOH both at mRNA levels (**Figure 4C**) and protein levels (**Figure 4D**), demonstrating that the MVA pathway is involved in the regulation of Atrogin-1 expression.

**Figure 4.**
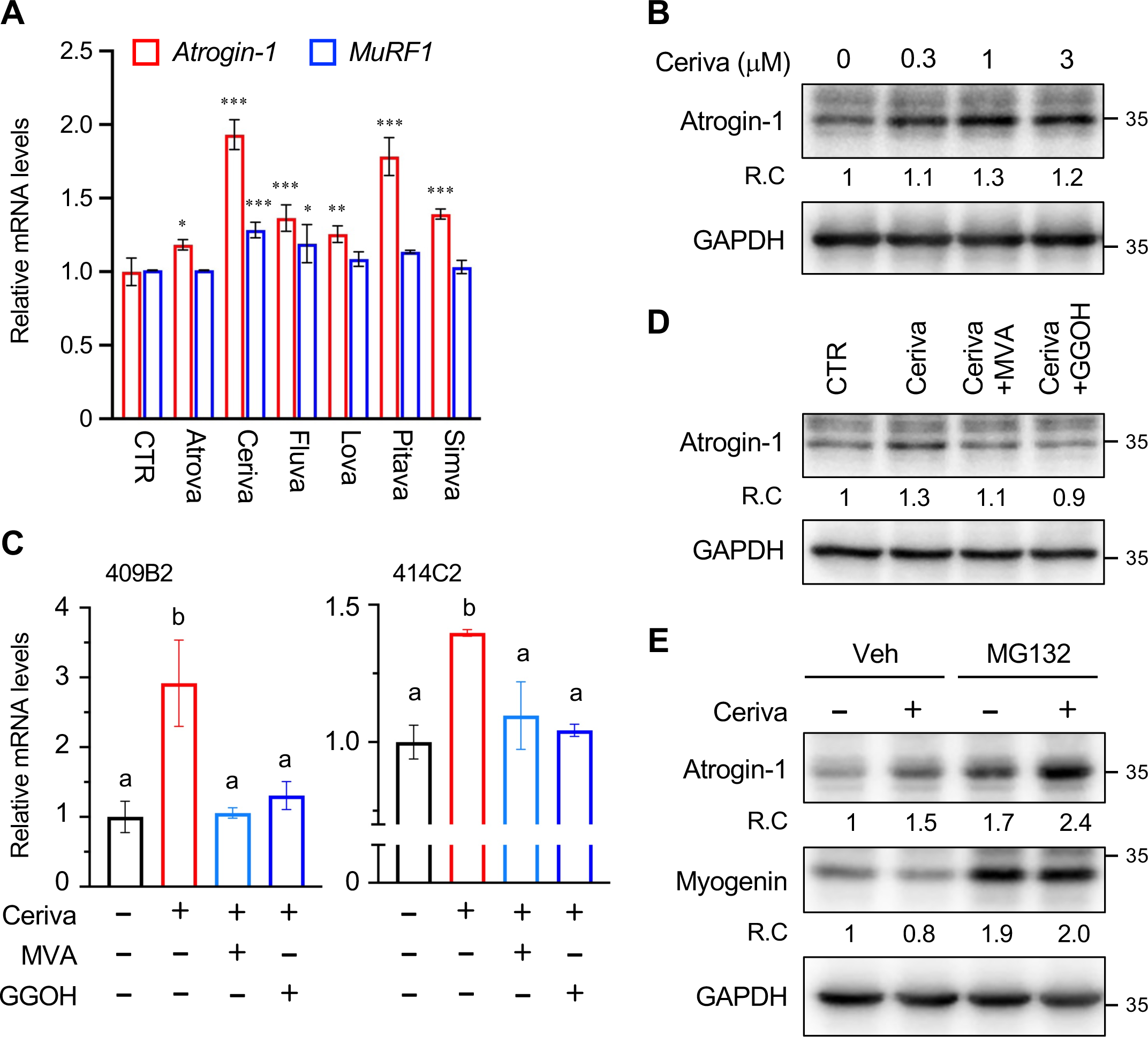
Statins promote Atrogin-1 expression and its substrate degradation in hiPSC-MCs. (A) Effects of different statins on *ATROGIN1* and *MURF1* gene expression. hiPSC-MCs (414C2) were incubated with the indicated statins (5 μM) for 16 h. mRNA levels of *ATROGIN1* and *MURF1* were measured by qRT-PCR. Data represent mean ± SD (n=3). Statistical analysis was performed by one-way ANOVA with Dunnett’s post hoc test (* p < 0.05, ** p < 0.01, *** p < 0.001). (B) Cerivastatin increases Atrogin-1 protein expression. hiPSC-MCs (414C2) were incubated with cerivastatin at the indicated concentrations for 16 h. Atrogin-1 expression was assessed by immunoblotting. GAPDH was used as an internal control. Relative changes (R.C.) of Atrogin-1 expression are shown at the bottom of the Atrogin-1 blot. (C, D) Statin-induced Atrogin-1 expression is reversed by MVA and GGOH. hiPSC-MCs (409B2 and 414C2) were incubated with cerivastatin (5 μM) in the presence or absence of MVA (200 μM) or GGOH (100 μM) for 16 h. Atrogin-1 mRNA levels (C) and protein levels (D) were assessed by qRT-PCR and immunoblotting, respectively. Data represent mean ± SD (n=3). Statistical analysis was performed by one-way ANOVA with Tukey’s post hoc test (C). Different letters denote statistical differences (p < 0.01). Relative changes (R.C.) of Atrogin-1 protein levels are shown at the bottom of the Atrogin-1 blot (D). (E) Cerivastatin induces proteasomal degradation of Myogenin. hiPSC-MCs (414C2) were incubated with 5 μM cerivastatin for 16 h. MG132 (20 μM) was added 3 h before harvest. Protein levels of Atrogin-1 and Myogenin were assessed by immunoblotting. GAPDH served as a loading control. Relative changes (R.C.) of Atrogin-1 and Myogenin expression are shown at the bottom of each blot.

To determine the functional significance of statin-induced Atrogin-1 expression, we examined whether cerivastatin promotes the proteasomal degradation of Myogenin (also known as MyoG), an Atrogin-1 substrate that is crucial for regulating muscle mass and function (29). Treating hiPSC-MCs with cerivastatin increased Atrogin-1 but decreased Myogenin protein levels (**Figure 4E**). Cerivastatin did not affect Myogenin mRNA levels (**Figure S3**), suggesting it promotes the degradation of Myogenin protein. To directly assess the proteasomal degradation of Myogenin, hiPSC-MCs were treated with cerivastatin in the presence or absence of MG132, a proteasome inhibitor. MG132 markedly increased Myogenin protein levels, indicating that Myogenin is degraded by the proteasome pathway. Furthermore, in the presence of MG132, Myogenin protein levels were unchanged by cerivastatin, suggesting that Atrogin-1 mediates the proteasomal degradation of Myogenin. Figure 4E also implies that Atrogin-1 is degraded by the proteasome pathway through an E3 ubiquitin ligase yet uncharacterized. Together, these results support our view that disrupting the MVA pathway upregulates Atrogin-1 expression, which accelerates the proteasomal degradation of its substrates and contributes to statin myotoxicity.

### Statins attenuate Akt signaling in hiPSC-MCs

The above results indicate that blocking the MVA pathway impairs the balance between protein synthesis and degradation in skeletal muscle. Since Akt signaling plays diverse important roles in regulating protein synthesis and degradation, we examined whether statins inhibits Akt signaling in hiPSC-MCs. The results showed that cerivastatin diminishes Akt phosphorylation in a concentration-dependent manner in hiPSC-MCs (**Figures 5A, S4A**), corroborating previous reports showing Akt inhibition by statins in murine myotubes (30–32). We next examined whether the inhibition of Akt by cerivastatin depends on the MVA pathway. The addition of MVA or GGOH restored the phosphorylation of Akt (**Figures 5B, S4B**), indicating that the Akt inactivation by cerivastatin results from the blockage of the MVA pathway. To further confirm these results, hiPSC-MCs were treated or untreated with cerivastatin and then stimulated with insulin-like growth factor-1 (IGF-1) to boost Akt signaling. The results showed that in hiPSC-MCs treated with cerivastatin, Akt phosphorylation by IGF-1 is diminished compared to untreated myocytes (**Figure S5**). The modest reduction in Atrogin-1 expression in response to the IGF-1 stimulation was also observed as reported (33).

**Figure 5.**
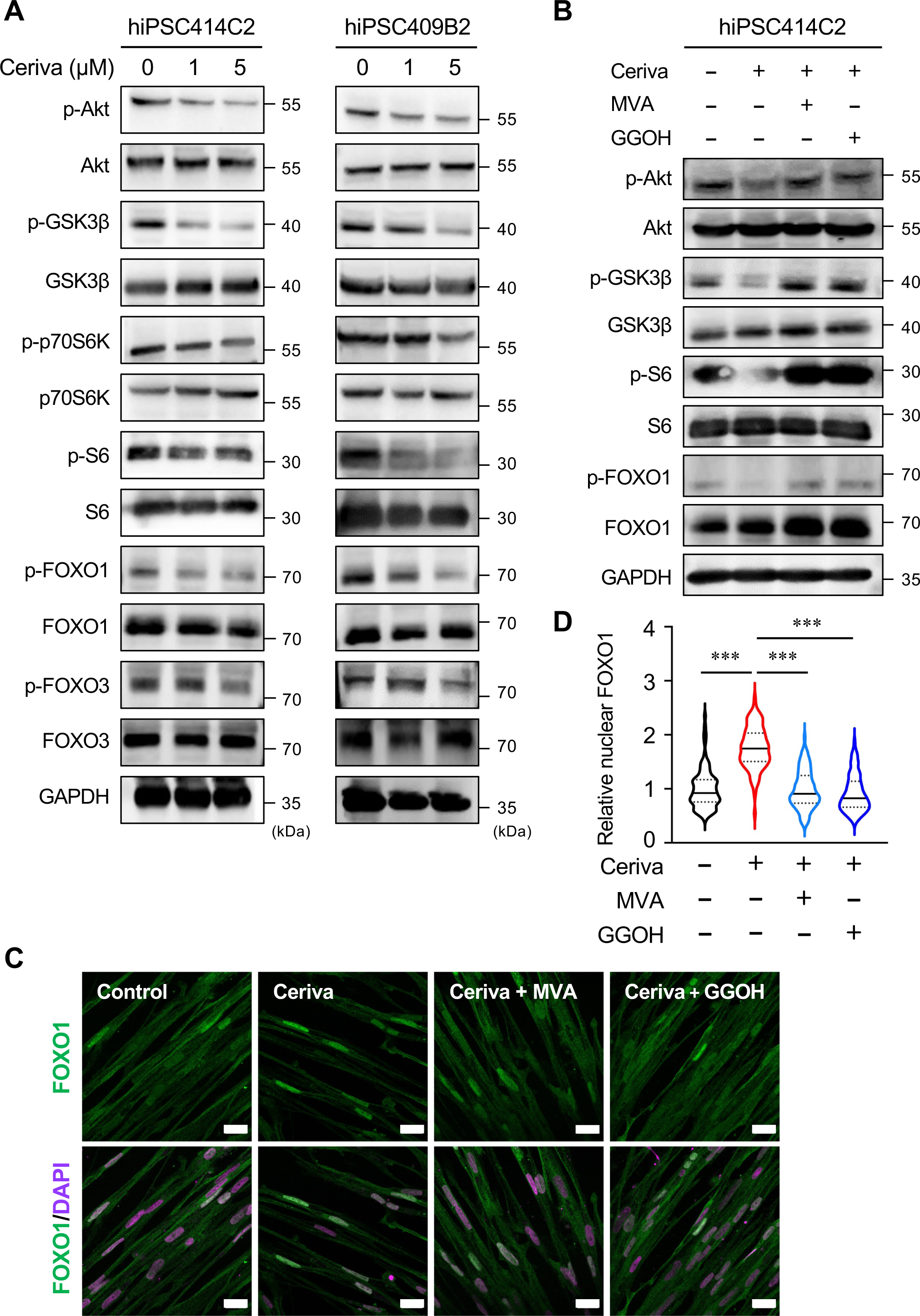
Blocking the MVA pathway disrupts the Akt signaling in hiPSC-MCs. (A) Cerivastatin suppresses Akt signaling. hiPSC-MCs (414C2 and 409B2) were treated with different concentrations of cerivastatin for 16 h. Cell lysate was subjected to immunoblot analysis to detect the indicated proteins. Relative changes in phosphorylation of each protein are shown in Figure S4A. (B) Effect of MVA and GGOH supplementation on Akt signaling. hiPSC-MCs (414C2) were treated with 5 μM cerivastatin in the presence or absence of MVA (200 μM) or GGOH (100 μM) for 16 h. Cell lysate was subjected to immunoblot analysis as above. Relative changes in phosphorylation of each protein are shown in Figure S4B. (C, D) Effect of the MVA pathway on FOXO1 localization. hiPSC-MCs (414C2) were treated with 5 μM cerivastatin in the presence or absence of MVA (200 μM) or GGOH (100 μM) for 16 h. Cells were stained for FOXO1 (green) and the nuclei (magenta). Scale bar, 20 μm (C). Nuclear FOXO1 intensities were shown in panel D. The results are mean ± SD (n=100). Statistical analysis was performed by one-way ANOVA with Tukey’s post hoc test (*** p < 0.0001).

To assess the effect of cerivastatin on effector molecules downstream of Akt signaling, we examined phosphorylation levels of GSK3β and FOXOs, direct Akt substrates, and p70S6K and S6, substrates of mTORC1 and p70S6K, respectively; these proteins play roles in the regulation of proteostasis (21–23). The results showed that phosphorylation of GSK3β, FOXO1, FOXO3, p70S6K, and S6 is all reduced by cerivastatin, but the effect on FOXO3 was subtle compared to FOXO1 (**Figures 5A, S4A**). The reduction in FOXO1 phosphorylation by cerivastatin was observed in hiPSC-MCs treated with IGF-1 (**Figure S5**). The addition of MVA or GGOH restored the reduction in phosphorylation of these proteins by cerivastatin (**Figures 5B, S4B**), confirming an important role of the MVA pathway in the Akt signaling cascade.

To further validate these results, the cellular distribution of FOXO1 was assessed by immunofluorescence confocal microscopy. The results showed that cerivastatin treatment markedly increases the nuclear accumulation of FOXO1 in hiPSC-MCs (**Figure 5C, D**). Moreover, in line with the above results, MVA and GGOH reversed the statin-dependent increase in nuclear localization of FOXO1. Cerivastatin had almost no effect on *FOXO1* and *FOXO3* mRNA levels in hiPSC-MCs while it upregulated the expression of *FOXO4* and *FOXO6* in a MVA pawathy-dependent manner (**Figure S6**). Collectively, these results indicate that statins inhibit the Akt–mTORC1 signaling, but activate the FOXO1 axis in hiPSC-MCs, which leads to the disruption of proteostasis in human myocytes.

### FOXO1 blockage prevents statin myotoxicity in hiPSC-MCs

To elucidate the pathophysiological significance of the FOXO1–Atrogin-1 axis, we sought to determine whether FOXO1 blockage can protect hiPSC-MCs from statin myotoxicity. We first examined the effect of the FOXO1 inhibitor AS1842856 on the expression of Atrogin-1. hiPSC-MCs were treated or untreated with cerivastatin in the presence or absence of AS1842856 for 16 h, and mRNA and protein levels of Atrogin-1 were assessed. The results showed that both in basal and cerivastatin-treated conditions, the FOXO1 inhibitor suppresses Atrogin-1 expression by approximately 50 % both at mRNA and protein levels (**Figure 6A, B**). The increase in Atrogin-1 expression by cerivastatin was canceled by FOXO1 blockage. These results suggest that FOXO1 plays a more dominant role in the upregulation of Atrogin-1 expression by cerivastatin in human myocytes.

**Figure 6.**
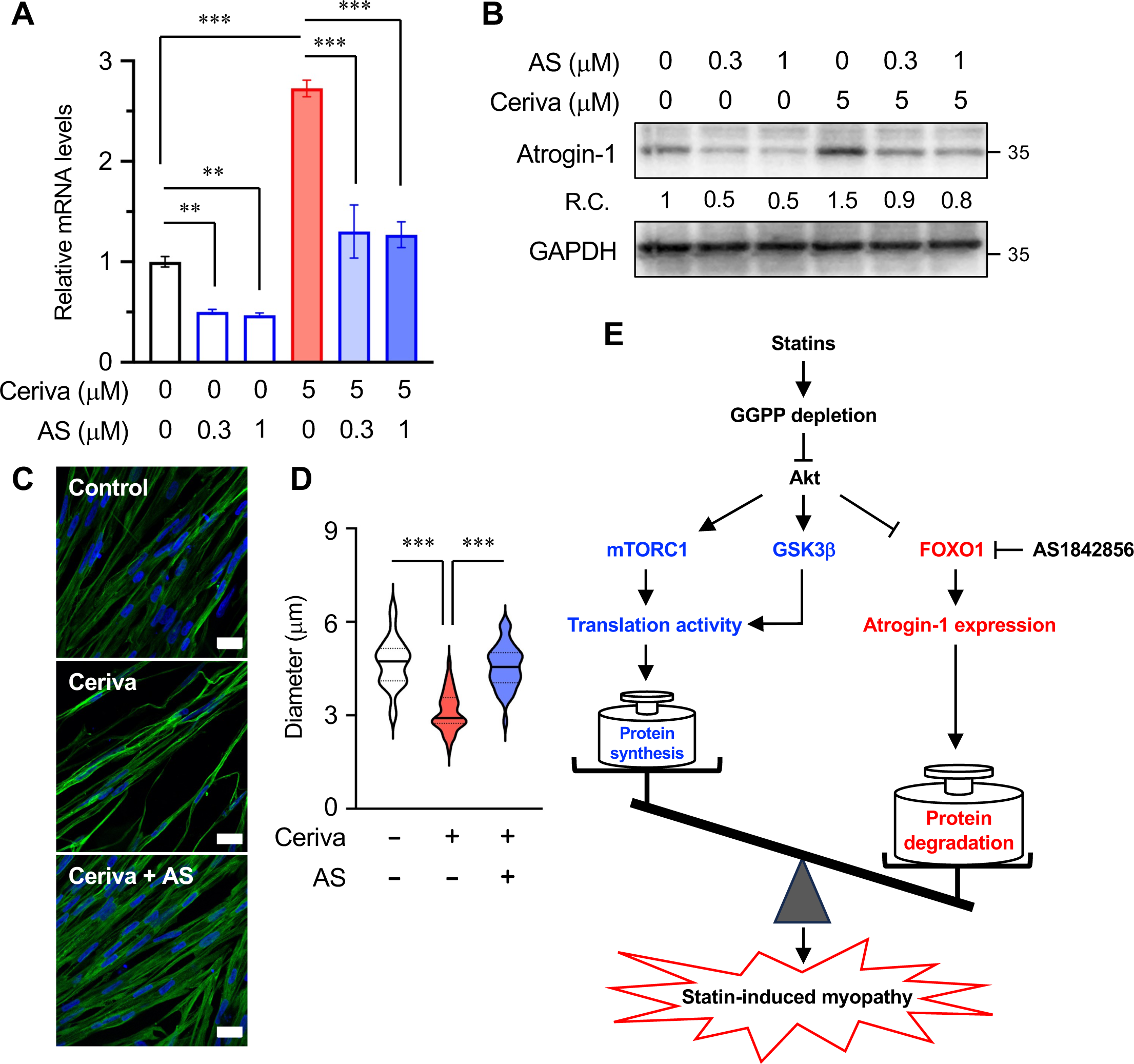
FOXO1 blockage prevents statin-induced atrogin-1 expression and myotoxicity in hiPSC-MCs. (A, B) FOXO1 inhibitor suppresses statin-induced expression of Atrogin-1. hiPSC-MCs (414C2) were treated with cerivastatin in the presence or absence of AS1842856 for 16 h. Atrogin-1 expression was assessed by qRT-PCR (A) and immunoblotting (B). Data are mean ± SD (n=3). Statistical analysis was performed by one-way ANOVA with Tukey’s post hoc test (** p < 0.01, *** p < 0.001). Relative changes in Atrogin-1 protein expression are shown below the Atrogin-1 blot. (C, D) Effect of FOXO1 inhibitor on statin-induced atrophy. hiPSC-MCs (414C2) were incubated with or without cerivastatin (5 μM) and AS1842856 (1 μM) for 16 h. hiPSC-MCs were stained with anti-MHC antibody (green) and DAPI (blue) (C). Scale bar; 20 μm. Diameters of MHC-positive myocytes were measured and expressed in violin plots (D). The results are mean ± SD (n=35). Statistical analysis was performed by one-way ANOVA with Tukey’s post hoc test (*** p < 0.0001). (E) Working model for SIM. Statin-dependent reduction in GGPP production suppresses Akt signaling and disrupts proteostasis in skeletal muscle. See text for more details.

Finally, we assessed the effect of FOXO1 blockage on statin-induced atrophy. hiPSC-MCs were incubated with cerivastatin in the presence or absence of AS1842856, and the diameters of MHC-positive myocytes were measured. The results showed that pharmacological blockage of FOXO1 prevents statin-induced atrophy (**Figure 6C, D**). Collectively, these results indicate that FOXO1 is a critical mediator of myotoxicity and suggest that blocking FOXO1 is sufficient to protect myocytes from SIM.

## Discussion

Human iPSCs have been utilized for modeling human diseases, which has been facilitating the elucidation of disease mechanisms and the development of therapeutic strategies (24, 25). This work was designed to establish an hiPSC-based SIM model for better understanding mechanisms of and validating therapeutic targets for SIM. Although the mechanisms of SIM have been extensively investigated, most studies used rodent C2C12 and L6 myocytes as in vitro models. Consequently, much less is known about how SIM is developed in human skeletal muscle. The present work is the first attempt to apply hiPSCs to study SIM, and our hiPSC-based model recapitulates SIM observed in the patients. Thus our hiPSC-based SIM model will provide important insights into the mechanisms of SIM and the development of therapeutic strategies for reducing statin intolerance.

Clinical and genetic studies demonstrate that disrupting the MVA pathway causes muscle weakness and damage, implying that the MVA pathway provides an important metabolite(s) for maintaining skeletal muscle homeostasis. Consistent with other studies in murine models (18), we showed that GGPP, not cholesterol or intermediate sterols, is an essential MVA-derived metabolite to prevent SIM. The requirement of GGPP for skeletal muscle is supported by evidence that squalene synthase or squalene epoxidase inhibitors do not have myotoxic effects (34, 35). Together, all these results indicate that the isoprenoid GGPP plays a crucial role in skeletal muscle homeostasis.

Skeletal muscle mass is strictly regulated by the rate of protein synthesis and degradation (22). Excessive proteolysis over protein synthesis results in muscle atrophy and a reduction of muscle strength, which are hallmarks of myopathy. For these reasons, we focused on proteostasis in myocytes in this work. We showed that cerivastatin and other statins inhibit de novo protein synthesis but increase protein degradation by upregulating the expression of Atrogin-1, a skeletal muscle-specific E3 ubiquitin ligase, in hiPSC-MCs, thereby disrupting proteostasis (**Figure 6E**). The statin-dependent increase in Atrogin-1 expression is consistent with clinical evidence reporting an increase in its expression in skeletal muscle biopsies from patients with SIM (36). Evidence shows that the risk of SIM increases with age (4, 37). In addition, it is suggested that aging impairs muscle proteostasis (28, 38). Accordingly, it is conceivable that statins augment the impairment of muscle proteostasis in aged patients, which leads to an increase in SIM risk in the elderly. This hypothesis needs further investigation.

How do statins disrupt muscle proteostasis? The protein kinase Akt plays diverse important roles in cell survival, proliferation, growth, and metabolism, and its downstream effectors regulate both protein synthesis and degradation (21). Inactivation of Akt by statins has been reported in murine C2C12 myocytes and skeletal muscle tissues of statin-treated rodents (30–32). Consistent with these reports, Akt was suppressed in hiPSC-MCs treated with cerivastatin, and MVA and GGOH at least partly reversed the effects. Furthermore, we showed that cerivastatin inhibits both mTORC1 and GSK3β, two major effectors that regulate protein translation, in a manner dependent on the MVA pathway, which explains the decrease in protein synthesis rate in statin-treated hiPSC-MCs. On the other hand, our results also revealed that blocking the MVA pathway inhibits the phosphorylation of FOXO1 and promotes its nuclear accumulation, thereby upregulating the expression of Atrogin-1. In addition, we showed that pharmacological blockage of FOXO1 attenuates statin-induced Atrogin-1 expression and cancels statin-induced myotoxicity. Collectively, the present study using a hiPSC-based SIM model suggests that blocking FOXO1 is sufficient to prevent statin myotoxicity. In addition, our findings suggest that Atorogin-1-dependent proteolysis plays a more dominant role than the reduction in protein synthesis in developing SIM. This view is supported by *Atrogin-1* null mice; myocytes from these mice exhibit statin resistance (36).

The importance of FOXO1 in preventing muscle atrophy has been reported in other settings; skeletal muscle-specific FOXO1 transgenic mice represent atrophic phenotype (39) whereas skeletal muscle-specific FOXO1 knockout mice are resistant to muscle wasting in chronic kidney disease (40). Recently, we have reported that blocking FOXO1 attenuates myostatin- and GDF11-dependent muscle atrophy in hiPSC-MCs (41). It is therefore conceivable that FOXO1 is a potential molecular target for preventing muscle wasting and myotoxicity in several contexts, including SIM and chronic kidney disease. Intriguingly, inhibition of FOXO1 also improves metabolism and is considered a therapeutic target for insulin resistance, diabetes, and non-alcoholic fatty liver disease (42, 43), which are often associated with elevated LDL-C. Therefore, statin therapy in combination with FOXO1 blockage could be a potential strategy for reducing the risk of cardiovascular disease with the prevention of SAMS.

In summary, this work highlights the physiological importance of the MVA pathway in maintaining skeletal muscle homeostasis. Our hiPSC-based SIM model suggests that statins disrupt Akt signaling and thus impair skeletal muscle proteostasis in humans; they not only suppress protein synthesis but also promote protein degradation through the FOXO1-Atrogin-1 axis, thereby accelerating myotoxicity. Moreover, the present study identifies FOXO1 as a potential target to prevent SIM.

## Experimental Procedures

### Materials

Statins were purchased from commercial sources as follows: atorvastatin (012–23901), fluvastatin (068–06641), lovastatin (125–04581), pitavastatin (163–24861), pravastatin (162–19821), and simvastatin (196–17801) from Fujifilm-Wako; cerivastatin (SML0005) and rosuvastatin (SML1264) from Sigma-Aldrich. Other chemicals were obtained as follows: mevalonic acid (90469), geraniol (163333), farnesol (277541), geranylgeraniol (G3278), cholesterol (C8667), and IGF-1 (SRP3069) from Sigma-Aldrich; cycloheximide (037–20991), squalene (194–09732), SB431542 (031–24291), and Y-27632 (036–24023) from Fujifilm-Wako; doxycycline (Dox) (D5897) from LKT laboratories; and AS1842856 (16761), farnesyl pyrophosphate (63250), geranyl pyrophosphate (63320), and geranygeranyl pyrophosphate (63330) from Cayman. Antibodies were obtained from the following sources: Anti-Akt (4691 or 9272; 1:1,000 dilution for immunoblot, IB), anti-phospho-Akt (Ser473) (4060; 1:2,000 dilution for IB), anti-FOXO1 (2880; 1:1,000 for IB; 1:100 for immunofluorescence, IF), anti-phospho-FOXO1 (Ser256) (9461; 1:1,000 for IB), anti-FOXO3a (12829; 1:1,000 for IB; 1:100 for IF), anti-FOXO3a (Ser253) (13129; 1:1,000 for IB), anti-GAPDH (2118; 1:1,000 for IB), anti-GSK3β (9315; 1:1,000 for IB), anti-phospho-GSK3β (Ser6) (9336; 1:1,000 for IB), anti-p70S6K (2708; 1:1,000 for IB), anti-phospho-p70S6K (Thr389) (9234; 1:1,000 for IB), anti-4E-BP1 (9644; 1:1,000 for IB), anti-phospho-4E-BP1 (Thr37/46) (2855; 1:1,000 for IB), anti-S6 (2217; 1:1,000 for IB), anti-phospho-S6 (Ser240/244) (2215; 1:1,000 for IB), HRP-linked anti-rabbit IgG (7074; 1:5,000 for IB), and HRP-linked anti-mouse IgG (7076; 1:5,000 for IB) from Cell Signaling Technology; anti-Fbx32/Atrogin-1 (ab168372; 1:1,000 for IB) from Abcam; anti-MuRF1 (sc-398608; 1:1,000 for IB) and anti-myogenin (sc-52903; 1:1,000 for IB) from Santa Cruz Biotechnology; anti-myosin heavy chain (MAB4470; 1:100 for IF) from R&D Systems; anti-puromycin antibody clone 12D10 (MABE343; 1:1,000 for WB) from Sigma-Aldrich; anti-SREBP-2 monoclonal antibody (clone 7D4) hybridoma supernatant and anti-HMGCR monoclonal antibody (clone A-9) hybridoma supernatant from Dr. Ta-Yuan Chang (Geisel School of Medicine at Dartmouth, NH).

### Cell culture

Undifferentiated hiPSCs (414C2^tet-MyoD^ and 409B2^tet-MyoD^) were maintained in a 6-well plate coated with iMatrix-511 silk (Matrixome) in StemFit AK02N (Ajinomoto) as described (44). The hiPSCs were differentiated into myocytes as described (41, 45). Briefly, the hiPSCs were seeded into a 6-well plate coated with Matrigel (356231, Corning) at a density of 3 × 10^5^ cells/well for 409B2^tet-MyoD^ or 4 × 10^5^ cells/well for 414C2^tet-MyoD^ and grown overnight in StemFit medium containing 10 μM Y-27632. On day 1, the medium was switched to Primate ES Cell Medium (Reprocell) containing Y-27632 (10 μM). On day 2, 1 μg/mL Dox was added to fresh Primate ES Cell Medium to induce MyoD expression, and cells were incubated overnight. On day 3, the medium was changed to αMEM supplemented with 5% KnockOut Serum Replacement (Gibco), 1 mg/mL Dox, 50 U/mL penicillin, and 50 mg/mL streptomycin (Nacalai Tesque), and the cells were further incubated for 2 to 3 days to be differentiated into myocytes. hiPSC-myocytes were incubated with DMEM supplemented with 5% horse serum (Gibco) overnight prior to experiments.

### Cell viability and cytotoxicity assays

hiPSCs were seeded into Matrigel-coated 24-well plates and differentiated as above. Cell viability and cytotoxicity were determined by Cell Counting Kit-8 (Donjindo) and Cytotoxicity LDH Assay Kit-WST (Donjido) according to the manufacturer’s protocols.

### Quantitative real-time PCR (qRT-PCR)

Total RNA was isolated using ISOGEN (Nippon Gene), and cDNA was synthesized from total RNA using a High-capacity cDNA Reverse Transcription Kit (Applied Biosystems). mRNA levels of a gene of interest were analyzed by qPCR using a specific primer set (**Table S1**) and FastStart Universal SYBR Green Master (Roche Applied Science). qRT-PCR was conducted using a StepOnePlus Real-Time PCR System (Applied Biosystems) or a Quant Studio Flex Real-Time PCR System (Applied Biosystems). 18S ribosomal RNA levels were used for normalization.

### Immunoblotting

Cells were lysed with urea buffer (8 M Urea, 50 mM Na-phosphate pH 8.0, 10 mM Tris-HCl pH 8.0, 100 mM NaCl) containing protease inhibitor cocktail (Nacalai Tesque) and phosphatase inhibitor cocktail (Sigma-Aldrich), vigorously vortexed for 30 min at 4 °C, and spun at 20,000 × *g* for 10 min at 4 °C as described (46). The supernatant was used as whole-cell lysate. Protein concentration was determined by BCA Protein Assay (Thermo Fisher Scientific). Equal amounts of cellular proteins were subjected to SDS-PAGE and immunoblot analysis according to a standard protocol. Signals were detected using FUSION Solo (Vilber), and band intensities were measured by Evolution Capt software (Vilber). GAPDH or α-tubulin (β-actin) was used as an internal control for normalization.

### Measurement of the rate of de novo protein synthesis

The protein synthesis rate was measured by the SUnSET method as described (26). Briefly, cells were incubated with 10 μg/mL of puromycin (Invivogen) for 60 min and then lysed with RIPA buffer (50 mM Tris-HCl pH 8.0, 150 mM NaCl, 1% (v/v) Triton X-100, 0.5% (w/v) deoxycholate, and 0.1% (w/v) SDS) with protease inhibitor cocktail (Nacalai Tesque). After protein concentration was determined using BCA Protein Assay, equal amounts of proteins were subjected to immunoblot analysis as above. The incorporation of puromycin into newly synthesized proteins was detected using anti-puromycin antibody and HRP-conjugated anti-mouse IgG antibody. Band intensities were quantified using ImageJ with α-tubulin as an internal control.

### Immunofluorescence staining and image analysis

hiPSCs were seeded into 35-mm film bottom dishes (FD10300, Matsunami) coated with Matrigel and differentiated into myocytes as above. After the treatment, hiPSC-MCs were fixed with 4% paraformaldehyde for 15 min, permeabilized with 0.1%Triton X-100 in PBS for 5 min, and blocked with 5% FBS in PBS for 1 h. Specimens were then treated with anti-MHC, anti-FOXO1, or anti-FOXO3 antibodies followed by incubation with Alexa Fluor 488-labeled goat anti-mouse IgG secondary antibody (1:800 dilution; A11029, Thermo Fisher Scientific) or Alexa Fluor 488-labeled goat anti-rabbit IgG secondary antibody (1:800 dilution; A11034, Thermo Fisher Scientific). Nuclei were stained with DAPI (Sigma-Aldrich). Specimens were mounted with ProLong Diamond Antifade Mountant (Thermo Fisher Scientific). Images were obtained using a Zeiss LSM800 confocal laser microscope equipped with a Plan-Apochromat 40×/1.40 Oil DIC M27 objective (Carl Zeiss) and processed with Zen software (Zeiss). The location of nuclei was identified by DAPI. Nuclear FOXO1 and FOXO3 levels were analyzed using Image J software. For measuring myocyte diameters, images were acquired by a fluorescence microscope BZ-X810 (KEYENCE) with a CFI S Plan Fluor ELWD 20×/0.45 objective (Nikon). Myocyte diameters were measured as described (33) using IC Measure (The Imaging Source). Briefly, diameters of MHC-positive myocytes were measured for each condition at three points, the thickest position and positions 50 μm apart from the thickest point, per myocyte as described (41).

### LC-MS/MS analysis

Cell extracts were prepared according to a method previously described (47) with modifications. Cells in 6-well plates were harvested in 2-propanol:100 mM NH_4_HCO_3_ (pH 7.8) (1:1 v/v) (100 μL/well) and collected into a 1.5 mL tube. Cells from three wells were pooled and used as a sample. Cells were then sonicated and homogenized with a 26-gauge needle for 10 strokes. Afterward, 2-propanol:100 mM NH_4_HCO_3_ (pH 7.8) (1:1 v/v) was added to the cell homogenate to be the final volume of 500 uL, and acetonitrile (500 μL/tube) was then added to the mixture for deproteinization. After placing on ice for 10 min, the mixture was centrifuged at 14,000 g for 10 min at 4 °C, and the supernatant was transferred to a glass tube. Subsequently, the supernatant was dried under a nitrogen stream at 40 °C, and the residues were dissolved in 80 μL of 1:1 (v/v) mixture of water and methanol. Samples were stored at −80 °C until use.

The LC-MS/MS analysis of nonsterol isoprenoids was conducted using a Nexera X3-LCMS-8060NX (Shimadzu) equipped with an ACCQ-TAG Ultra C18 column (1.7 μm, 100 mm x 2.1 mm I.D.; Waters) and a SIL-40C X3 autosampler at the Division of Analytical and Measuring Instruments, Shimadzu (Japan) as described (48) with modifications. The sample (10 μL/sample) was injected using an autosampler and separated by an ACCQ-TAG Ultra C18 column at a flow rate of 0.3 mL/min. Negative electrospray ionization (ESI) and multiple reaction monitoring MRM) were used for mass spectrometry. Detailed parameters for LC-MS/MS analysis are presented in **Table S2**. The retention time, MRM, and collision energy for each isoprenoid are summarized in **Table S3**. Standard curves for GPP, FPP, and GGPP were linear at the range of 0.005–1.0 μg/mL, 0.005–1.0 μg/mL, and 0.05–1.0 μg/mL, respectively. The amounts of isoprenoids were calculated with each standard curve and normalized to cellular protein determined by BCA assay.

### Statistical analysis

Results are presented as mean ± SD from at least three or more independent biological replicates. All experiments were repeated on at least two occasions with similar results. Statistical analyses were performed using unpaired Student’s *t*-test or one-way ANOVA with Turkey’s or Dunnett’s post hoc tests as specified in figure legends. *P* values less than 0.05 were considered statistically significant.

## Data availability

All data are contained within the manuscript.

## Supporting information

This article contains supporting information.

## Acknowledgments

We thank Dr. Ta-Yuan Chang (Geisel School of Medicine at Dartmouth, NH) for antibodies and the Division of Analytical and Measuring Instruments, Shimadzu for LC-MS/MS analysis.

## Conflict of interest

The authors declare that they have no conflict of interest with the contents of this article.

## Funding sources

This work was supported by AMED-CREST grant 21gm091008h (to Y.Y. and R.S.) from Japan Agency for Medical Research and Development, KAKENHI grants 19H02908 and 22H02281 (to Y.Y.) and 20H00408 (to R.S.) from the Japan Society for the Promotion of Science, the Nutrition and Food Science Fund of Japan Society of Nutrition and Food Science (to Y.Y.), and a research grant from the Tojuro Iijima Foundation for Food Science and Technology (to Y.Y.).

## Authors contributions

Y.Y. conceived and designed the study; X.Z., N.L. M.K., M.M., Y.Y. performed experiments; X.Z., N.L., M.K., R.S., and Y.Y. analyzed data; H.S. provided hiPSCs; Y.Y. and X.Z. wrote the manuscript; All authors read and approved the manuscript.

## Abbreviations

CVD: cardiovascular disease
ER: endoplasmic reticulum
FOH: farnesol
FPP: farnesyl pyrophosphate
GGOH: geranylgeraniol
GGPP: geranylgeranyl pyrophosphate
GOH: geraniol
hiPSCs: human induced pluripotent stem cells
HMG-CoA: 3-hydroxy-3-methylgrutaryl coenzyme A
HMGCR: HMG-CoA reductase
IPP: isopentenyl pyrophosphate
LDL: low-density lipoprotein
SAMS: statin-associated muscle symptoms
SIM: statin-induced myopathy
SREBP: sterol regulatory element-binding protein.

## Notes

### Competing Interest Statement

The authors have declared no competing interest.

