## Supplemental information for "Modeling statin-induced myopathy with human iPSCs reveals that impaired proteostasis underlies the myotoxicity and is targetable for the prevention"

Yoshio Yamauchi, Ph.D.

Department of Applied Biological Chemistry, Graduate School of Agricultural and Life Sciences, The University of Tokyo

1-1-1 Yayoi, Bunkyo-ku, Tokyo 113-8657, Japan

Supporting Tables: Table S1–S3

Supporting Figures: Figure S1–S5

Table S1. Primers used for qRT-PCR analysis of human genes

| Gene | Primer sequences |
|---|---|
| *FBXO32/ATROGIN1* | Fw: 5’-TGTTACCCAAGGAAAGAGCAGTATGGA-3’ |
| | Rv: 5’-ACGGAGCAGCTCTCTGGGTTATTG-3’ |
| *FOXO1* | Fw: 5’-ATGGATGGAGATACATTGGATTTTC-3’ |
| | Rv: 5’-CACTGTGTGGGAAGCTTTGGT-3’ |
| *FOXO3* | Fw: 5’-AGCCTAACCAGGGAAGTTTG-3’ |
| | Rv: 5’-GACTCACTCAAGCCCATGTT-3’ |
| *FOXO4* | Fw: 5’-ACGAGTGGATTGGTCCGTACT-3’ |
| | Rv: 5’-GTGGCGGATCGAGTTCTTC-3’ |
| *FOXO6* | Fw: 5’-ACCTCATCACCAAAGCCATC-3’ |
| | Rv: 5’-GCATCCACCACGAACTCTT-3’ |
| *HMGCS1* | Fw: 5’-GACTTGTGCATTCAAACATAGCAA-3’ |
| | Rv: 5’-GCTGTAGCAGGGAGTCTTGGTACT-3’ |
| *HMGCR* | Fw: 5’-TACCATGTCAGGGGTACGTC-3’ |
| | Rv: 5’-CAAGCCTAGAGACATAATCATC-3’ |
| *MYOG/MYOGENIN* | Fw: 5’-TGGGCGTGTAAGGTGTGTAA-3’ |
| | Rv: 5’-CGATGTACTGGATGGCACTG-3’ |
| *TRIM63/MURF1* | Fw: 5’-TGGGGGAGCCACCTTCCTCT-3’ |
| | Rv: 5’-ATGTTCTCAAAGCCCTGCTCTGTCT-3’ |
| *18S rRNA* | Fw: 5’-ACCGCAGCTAGGAATAATGGA-3’ |
| | Rv: 5’-GCCTCAGTTCCGAAAACCA-3’ |

Table S2. Conditions for LC-MS/MS analysis

| | |
|---|---|
| *LC parameters* | |
| Column | ACCQ-TAG Ultra C18 column (1.7 µm, 100 mm x 2.1 mm I.D) |
| Column temp. | 50 ºC |
| Mobile phase A | 10 mM ammonium carbonate with 0.1% ammonium hydroxide |
| Mobile phase B | Methanol with 0.1% ammonium hydroxide |
| Flow rate | 0.3 mL/min |
| *MS parameters* | |
| Ionization | negative electrospray ionization (ESI) |
| Interface voltage | - 4kV (-) |
| Ion focus voltage | - 4kV |
| Nebulizer gas flow rate | 2 L/min |
| Drying gas flow rate | 10 L/min |
| Heating gas flow rate | 10 L/min |
| Interface temp. | 350 ºC |
| Desolvation line temp. | 250 ºC |
| Heat block temp. | 400 ºC |
| Measurement mode | Multiple reaction monitoring (MRM) |

Table S3. Retention time, MRM transition, and collision energy for isoprenoid analysis.

| | RT (min) [1] | MRM transition (m/z) | CE (eV) [2] |
|---|---|---|---|
| GPP | 3.6 | 313.00 > 78.90 | 22.0 |
| FPP | 5.8 | 381.10 > 78.95 | 26.0 |
| GGPP | 7.2 | 449.15 > 78.90 | 26.0 |

[1] RT: Retention time

[2] CE: Collision energy

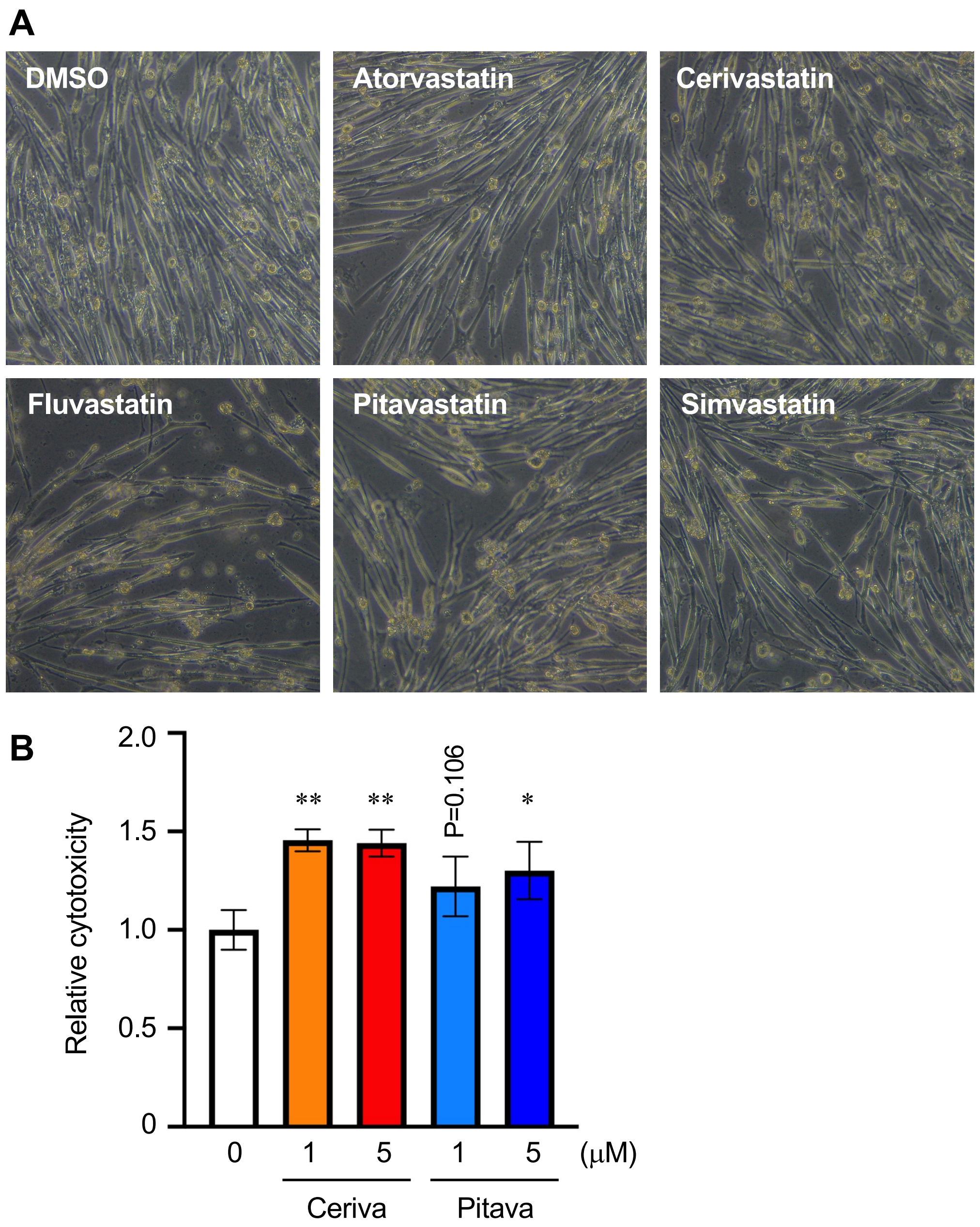


**Figure S1. Effect of different statins on myotoxicity in hiPSC-MCs.**

(A) hiPSC-MCs (414C2) were incubated with or without atorvastatin, cerivastatin, fluvastatin, pitavastatin, or simvastatin (5 μM each) for 18 h. Shown are typical images of hiPSC-MCs after the treatment.

(B) hiPSC-MCs (414C2) were incubated with or without cerivastatin or pitavastatin at the indicated concentrations for 18 h. Cytotoxicity was assayed as described in Materials and Methods. Results are means ± SD (n=3). Statistical analysis was performed by one-way ANOVA with Dunnett's post hoc test (* $p < 0.05$, ** $p < 0.01$).

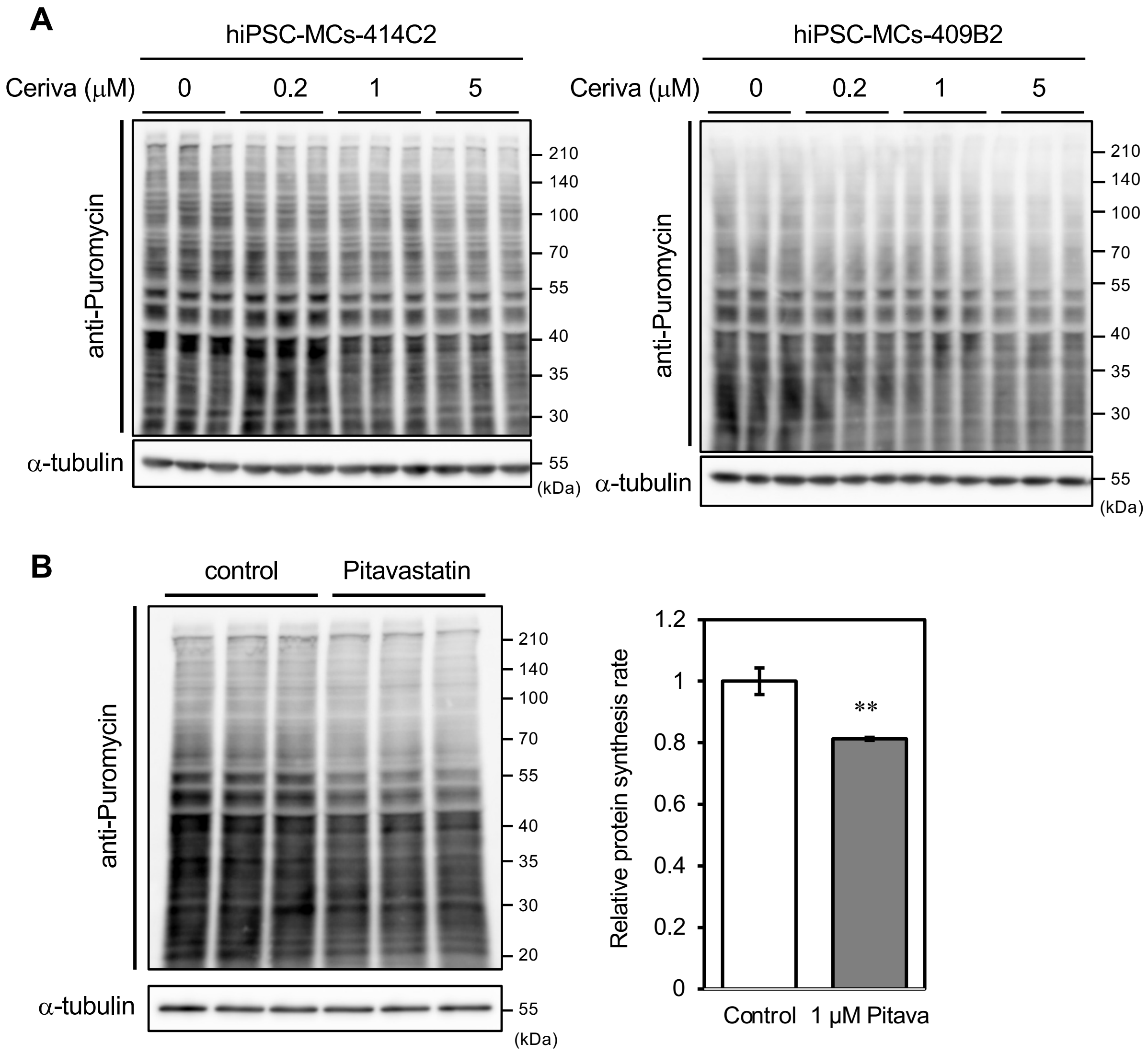


**Figure S2. Effect of statins on the global protein synthesis in hiPSC-MCs.**

(A) Immunoblot images used for the quantification in Figure 3A. hiPSC-MCs (414C2 and 409B2) were treated with cerivastatin at the indicated concentration for 16 h and then incubated with puromycin for 60 min. SUnSET assay was performed to evaluate the rate of protein synthesis as described in Materials and Methods. The quantification results are presented in Figure 3A.

(B) hiPSC-MCs (409B2) were incubated with or without pitavastatin (1 μM) for 16 h and then incubated with puromycin for 60 min. SUnSET assay was performed as above. Left: Immunoblot images, Right: Quantification result based on the immunoblot data. Results are means ± SD (n=3). Statistical analysis was performed by student's *t*-test (** $p < 0.01$).

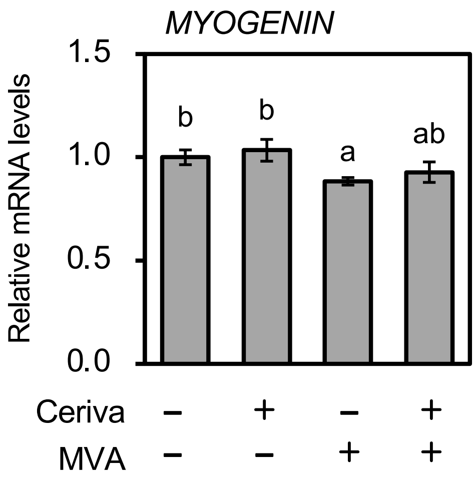


**Figure S3. Effect of cerivastatin on *MYOGENIN* mRNA expression.**

hiPSC-MCs (414C2) were incubated with 5 μM cerivastatin for 16 h in the presence or absence of MVA (0.2 mM). mRNA levels of *MYOGENIN* were examined. Data represent mean ± SD (n=3). Statistical analysis was performed by one-way ANOVA with Tukey's post hoc test. Different letters denote statistical differences ($p < 0.05$).

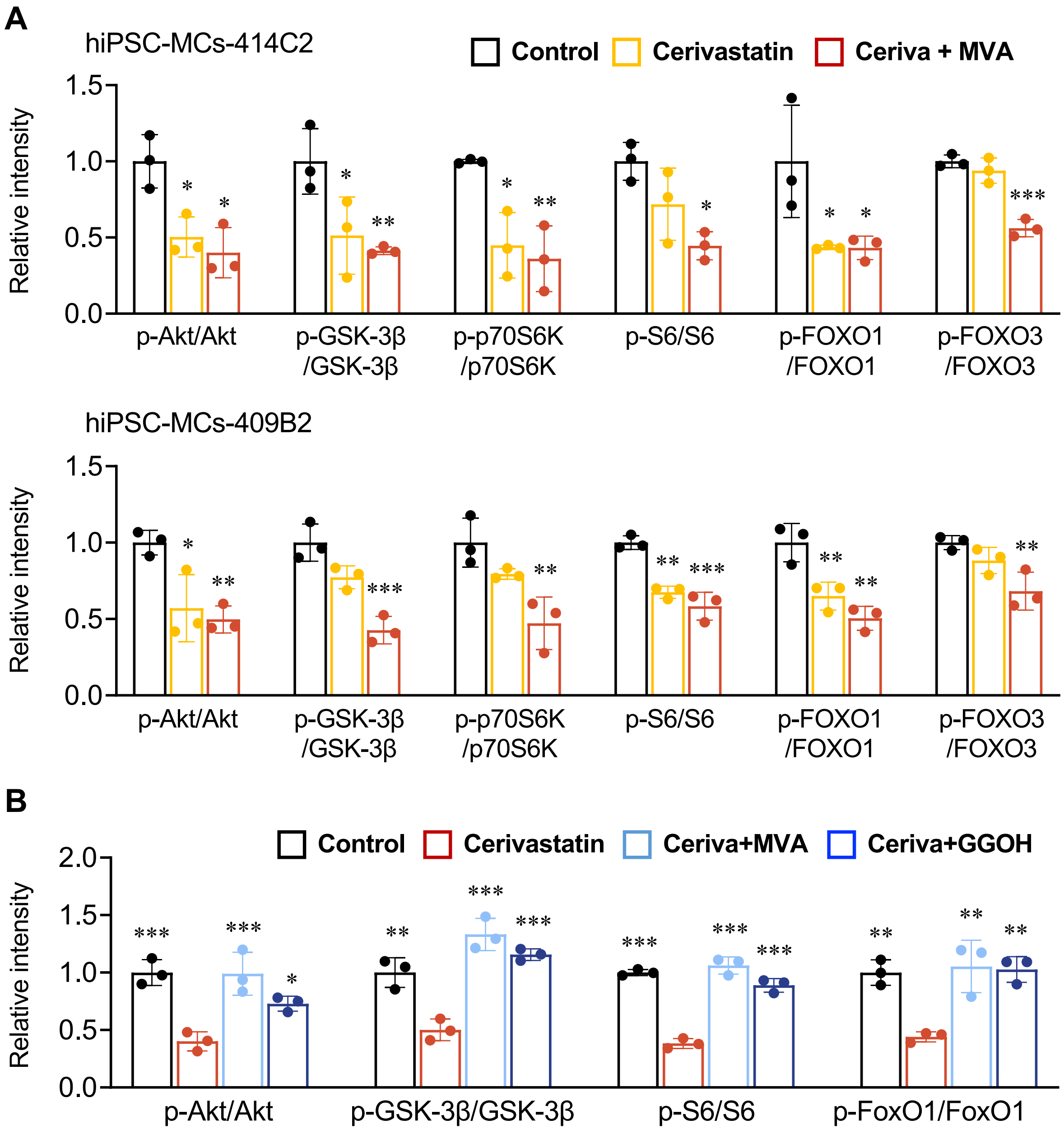


**Figure S4. Cerivastatin attenuates Akt signaling in hiPSC-MCs.**

(A) Quantification results of Figure 5A.

(B) Quantification results of Figure 5B.

Data are mean ± SD (n=3). Statistical analyses were performed by one-way ANOVA with Dunnett's post hoc test (* $p < 0.05$, ** $p < 0.01$).

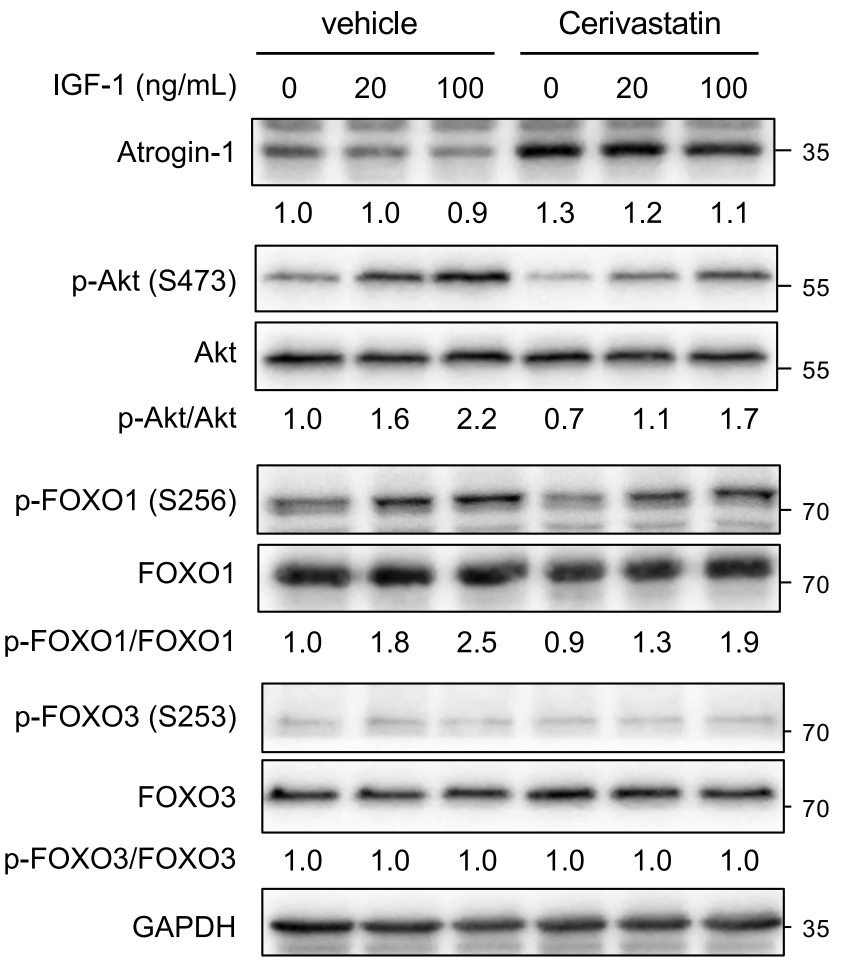


**Figure S5. Cerivastatin attenuates IGF-1-dependent Akt signaling in hiPSC-MCs.**

hiPSC-MCs (414C2) were incubated with or without cerivastatin (5 μM) for 16 h. Myocytes were then stimulated with 20 or 100 ng/mL of IGF-1 for 30 min in the presence or absence of cerivastatin. Cell lysate was subjected to immunoblotting to analyze the expression of Atrogin-1 and the phosphorylation of Akt, FOXO1, and FOXO3. Relative changes are shown at the bottom of each blot.

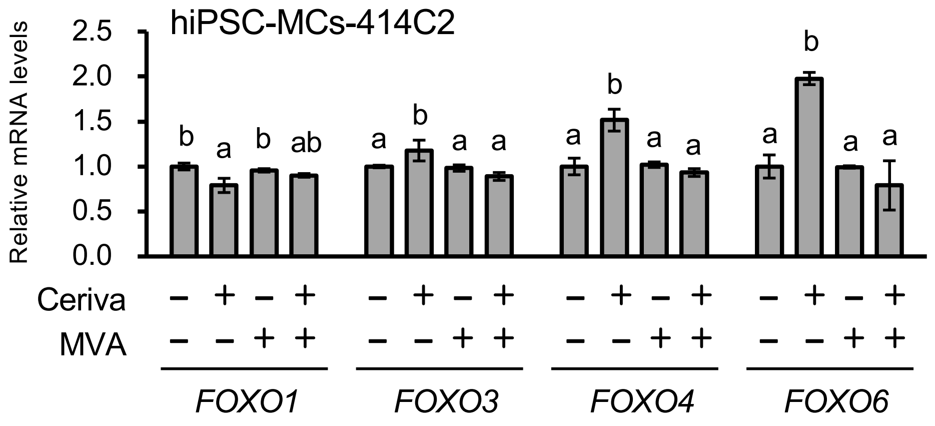


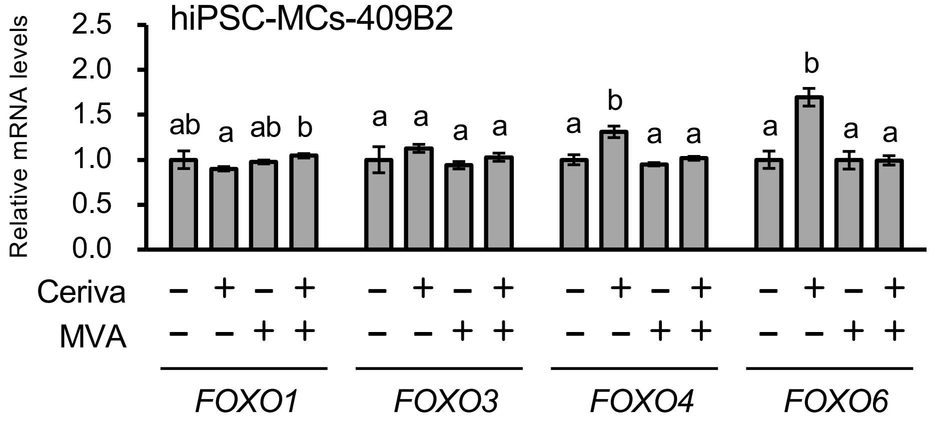


**Figure S6. Effects of the MVA pathway on mRNA levels of FOXOs.**

hiPSC-MCs (414C2 and 409B2) were incubated with or without cerivastatin (5 μM) and MVA (200 μM) for 16 h as indicated. mRNA levels of *FOXO1*, *FOXO3*, *FOXO4*, and *FOXO6* were analyzed by qRT-PCR. Results are means ± SD (n=3). Statistical analyses were performed by one-way ANOVA with Tukey's post hoc test. Different letters denote statistical differences ($p < 0.05$).
